## Supplementary materials for "A *Ctnnb1* enhancer transcriptionally regulates Wnt signaling dosage to balance homeostasis and tumorigenesis of intestinal epithelia"

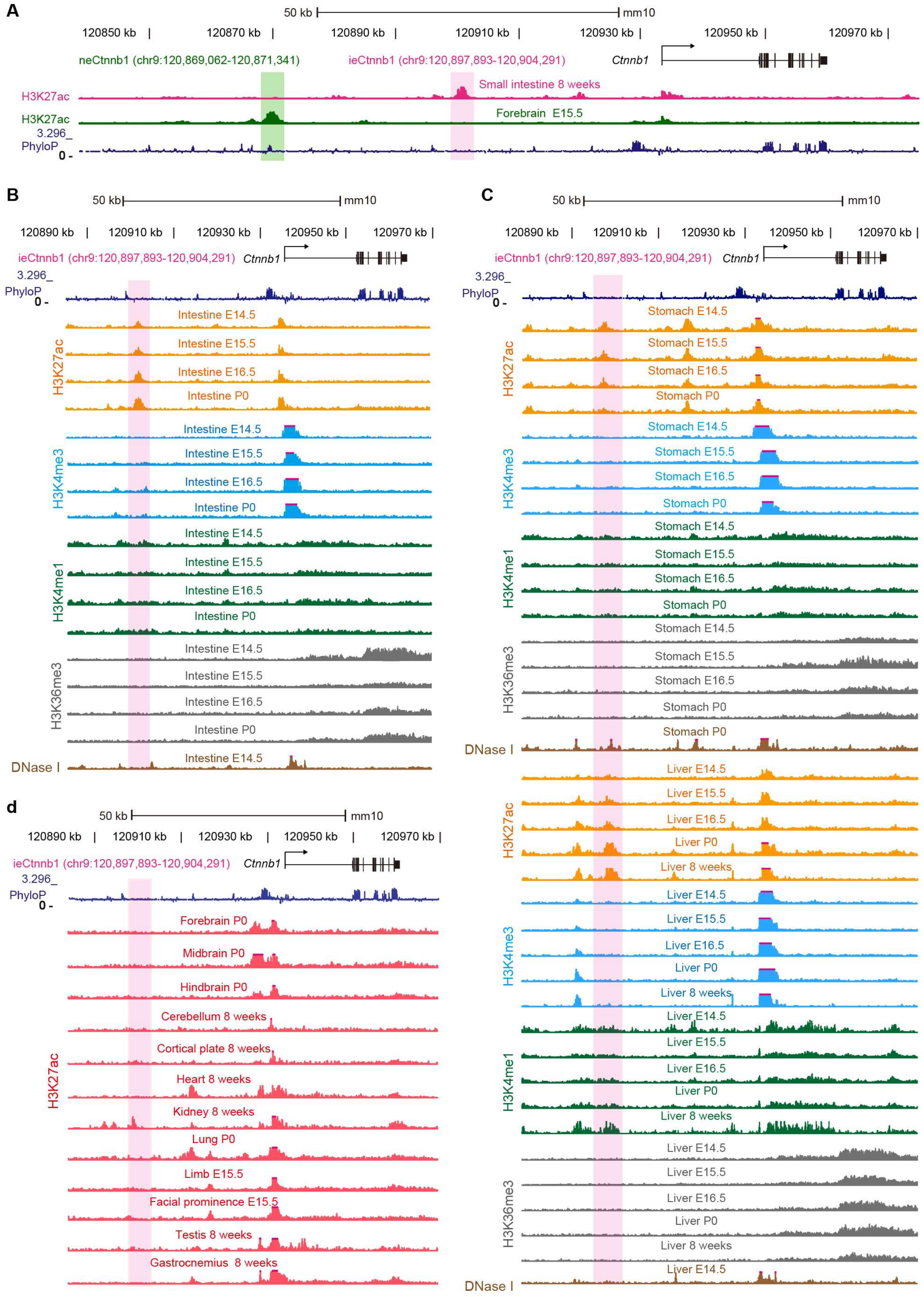


**Figure 1 - figure supplement 1. ieCtnnb1 is a putative intestinal enhancer upstream of *Ctnnb1*.** (A) Schematic representation of the upstream region of mouse *Ctnnb1* gene and locations of enhancer neCtnnb1 (green shading) and putative enhancer ieCtnnb1 (pink shading). Data were obtained from ENCODE. (B-C) Enrichment of indicated signals in developing intestine (B), stomach and liver (C) were shown. Data were obtained from ENCODE. (D) Enrichment of H3K27ac ChIP-seq signals in indicated mouse tissues. Data were obtained from ENCODE.

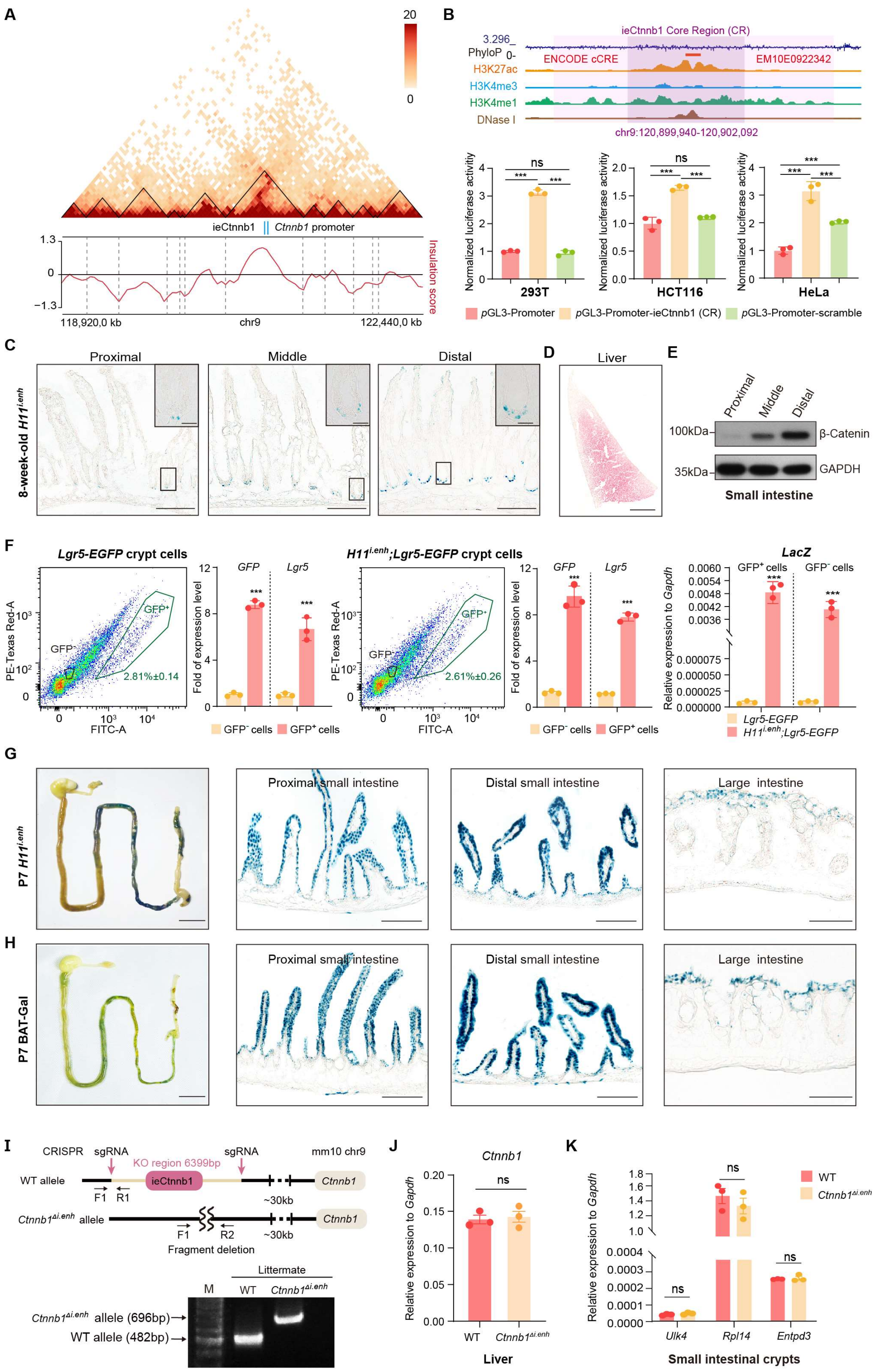


**Figure 1 - figure supplement 2. ieCtnnb1 is predominantly active in**

**developing intestine.** (A) Hi-C data of BALB/c mouse large intestine at indicated locus. Boundaries of TADs and locations of *Ctnnb1* promoter and ieCtnnb1 (blue bars) were marked. (B) Top: schematic representation of ieCtnnb1 core region (2,153 bp, dark purple shading). The location of an annotated ENCODE cCRE was indicated. Bottom: HEK293T, HCT116 and HeLa cells were transfected with indicated plasmids for 48 hours for luciferase reporter assay. (C) Representative images showing X-Gal staining of proximal, middle, and distal small intestinal sections of 8-week-old $H11^{i.enh}$ mice (blue). (D) Representative image showing X-Gal and eosin staining of the liver of 8-week-old $H11^{i.enh}$ mice (X-Gal: blue, eosin: red). (E) Immunoblotting of indicated proteins derived from proximal, middle, and distal small intestine tissues of WT C57/BL6 mice. (F) Flow cytometry assays to isolate GFP+ and $GFP^-$ cells from small intestinal crypts of *Lgr5*-EGFP (n = 3) and $Ctnnb1^{\Delta i.enh}$;*Lgr5*-EGFP (n = 3) mice. RT-qPCR showing relative mRNA levels of *GFP, Lgr5* and *LacZ* in aforementioned GFP+ and $GFP^-$ cells. (G-H) Representative images showing X-Gal staining of gastrointestinal tracts, and sections of proximal and distal small intestine, and large intestine of P7 $H11^{i.enh}$ (G) mice and P7 BAT-Gal mice (H). (I) Generation and genotyping of $Ctnnb1^{\Delta i.enh}$ mice. WT, wild-type; sgRNA, small guide RNA. (J) Relative mRNA levels of *Ctnnb1* in the liver of WT (n = 3) and $Ctnnb1^{\Delta i.enh}$ (n = 3) mice. (K) Relative mRNA levels of genes (*Ulk4, Rpl14* and *Entpd3)* located in the same TAD as *Ctnnb1* in small intestinal crypts of WT (n = 3) and $Ctnnb1^{\Delta i.enh}$ (n = 3) mice. Scale bars, 1 cm (whole mount in F and G), 100 μm (C, D, F and G), 10 μm (magnified insets in C). Quantification data are shown as means ± SEM. Statistical significance was determined using Two-way ANOVA analysis (B) and an unpaired two-tailed Student's *t*-test (F, J and K). *$P < 0.05$, **$P < 0.01$, ***$P < 0.001$, and ****$P < 0.0001$. ns, not significant.

A

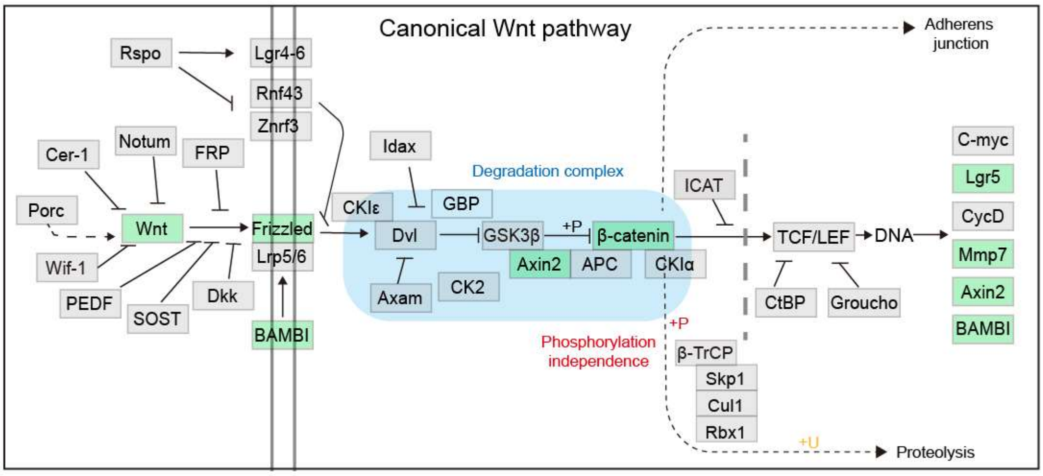


B

| Wnt signalling components | Gene | $Log_2$ Fold Change | Explanation | Reference |
|---|---|---|---|---|
| Wnt ligands | *Wnt3* | -1.16 | Decreased numbers of Paneth cells | Sato et al., 2011 (PMID21113151) |
| | *Wnt2b* | -0.55 | | |
| Wnt receptors | *Frizzled3* | -1.28 | Possible cellular adaptation upon reduced Wnt ligands | — |
| | *Frizzled9* | -0.60 | | |
| β-catenin | *Ctnnb1* | -0.27 | Knockout of ieCtnnb1 | — |
| Target genes | *Axin2* | -0.36 | Reduced β-catenin level in the nucleus | Jho et al., 2023 (PMID11809808) |
| | *Lgr5* | -0.44 | | Barker & Clevers, 2010 (PMID20417836) |
| | *Mmp7* | -0.73 | | Crawford, 1999 (PMID10362259) |
| | *Bambi* | -0.71 | | Sekiya, 2004 (PMID14660579) |

**Figure 1 - figure supplement 3. The list of Wnt signaling pathway components downregulated in *Ctnnb1*$^{\Delta i.enh}$ crypts.**

(A) A schematic diagram highlighting downregulated components of the Wnt signaling pathway in the crypts of the *Ctnnb1*$^{\Delta i.enh}$ small intestine. (B) A table showing Wnt signaling pathway components downregulated in *Ctnnb1*$^{\Delta i.enh}$ crypts.

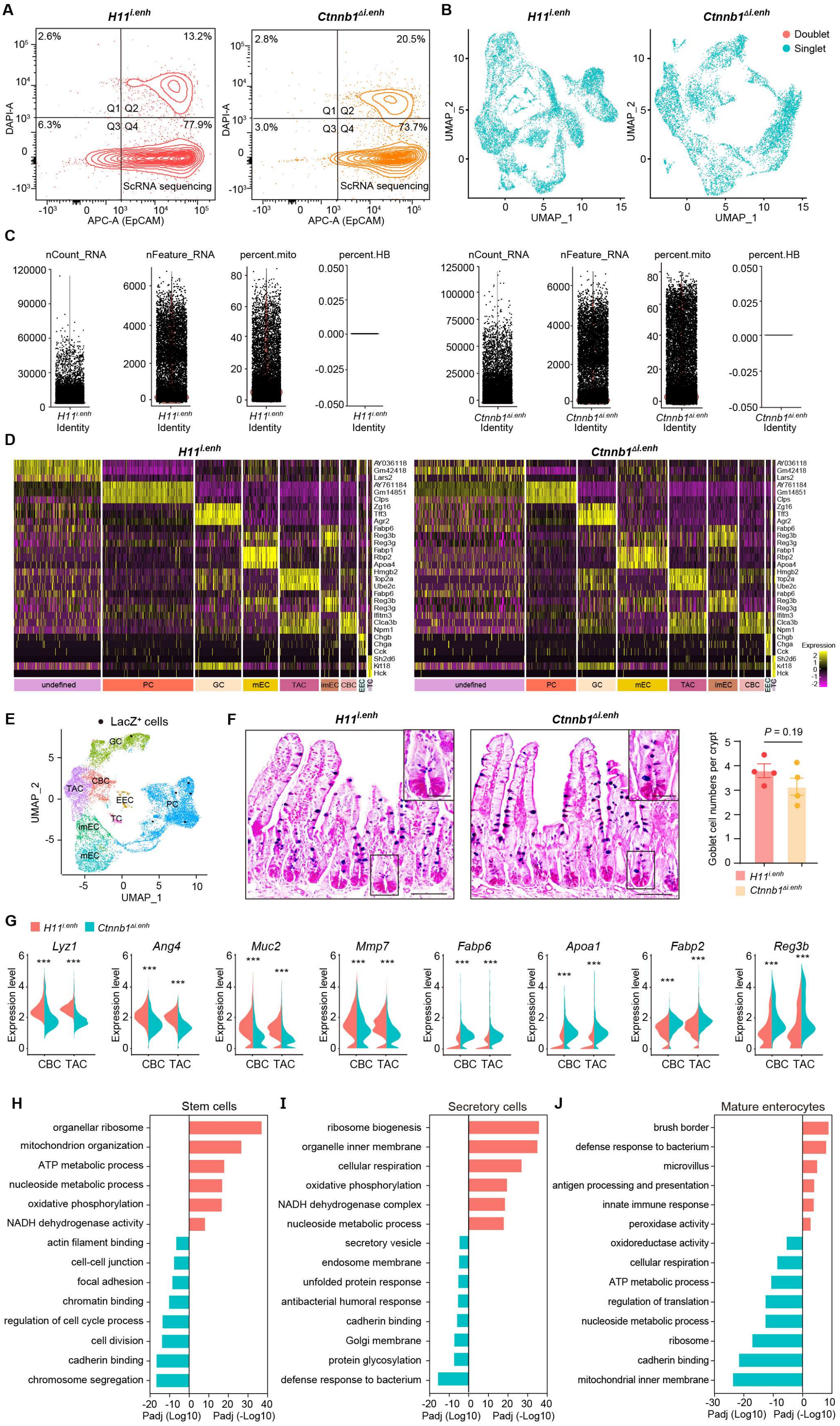

A
B
C
D
E
F
G
H
I
J
ScRNA sequencing
Doublet
Singlet
LacZ⁺ cells
P = 0.19
Goblet cell numbers per crypt
Stem cells
Secretory cells
Mature enterocytes
organellar ribosome
mitochondrion organization
ATP metabolic process
nucleoside metabolic process
oxidative phosphorylation
NADH dehydrogenase activity
actin filament binding
cell-cell junction
focal adhesion
chromatin binding
regulation of cell cycle process
cell division
cadherin binding
chromosome segregation
ribosome biogenesis
organelle inner membrane
cellular respiration
oxidative phosphorylation
NADH dehydrogenase complex
nucleoside metabolic process
secretory vesicle
endosome membrane
unfolded protein response
antibacterial humoral response
cadherin binding
Golgi membrane
protein glycosylation
defense response to bacterium
brush border
defense response to bacterium
microvillus
antigen processing and presentation
innate immune response
peroxidase activity
oxidoreductase activity
cellular respiration
ATP metabolic process
regulation of translation
nucleoside metabolic process
ribosome
cadherin binding
mitochondrial inner membrane
Padj (Log10)
Padj (-Log10)

**Figure 2 - figure supplement 1. Single-cell survey of small intestinal crypt cells upon ieCtnnb1 knockout.** (A) Flow cytometry assays for isolating EpCAM+DAPI- epithelial cells from $H11^{i.enh}$ (n=2) and $Ctnnb1^{\Delta i.enh}$ (n=2) small intestinal crypts. (B) Results of cell filtration using DoubletFinder. 1,294 doublets of 11,824 cells from $H11^{i.enh}$ crypts and 567 doublets of 8094 cells from $Ctnnb1^{\Delta i.enh}$ crypts were identified (blue dots: singlets, red dots: doublets). (C) Results of cell filtration using Seurat. Cells with more than 30% of reads derived from mitochondrial genes are considered dead and removed from the analysis. (D) Cell-type specific signatures. Heatmap showing relative expression levels (row-wise Z scores) of genes (rows) in cell-type-specific signatures. (E) UMAP visualizing LacZ+ single cells (black dots) of $H11^{i.enh}$ small intestinal crypts (n = 4 mice). (F) Alcian Blue PAS staining and quantification of goblet cells in the small intestine of $H11^{i.enh}$ (n = 6) and $Ctnnb1^{\Delta i.enh}$ (n = 6) mice. (G) Violin plots showing expressions of marker genes for secretory cells (*Lyz1, Ang4, Muc2, Mmp7*) and absorptive cells (*Fabp6, Apoa1, Fabp2, Reg3b*) in $H11^{i.enh}$ and $Ctnnb1^{\Delta i.enh}$ CBCs and TACs. (H-J) Gene ontology (GO) analysis of differential genes in stem cell lineage (H), secretory lineage (I), and mature enterocytes (J) of $H11^{i.enh}$ and $Ctnnb1^{\Delta i.enh}$ crypts. Scale bars, 50 μm (E), 10 μm (magnified views in E). Quantification data are shown as means ± SEM, statistical significance was determined using an unpaired two-tailed Student's *t*-test (F and G). **P* < 0.05, ***P* < 0.01, ****P* < 0.001, and *****P* < 0.0001. ns, not significant.

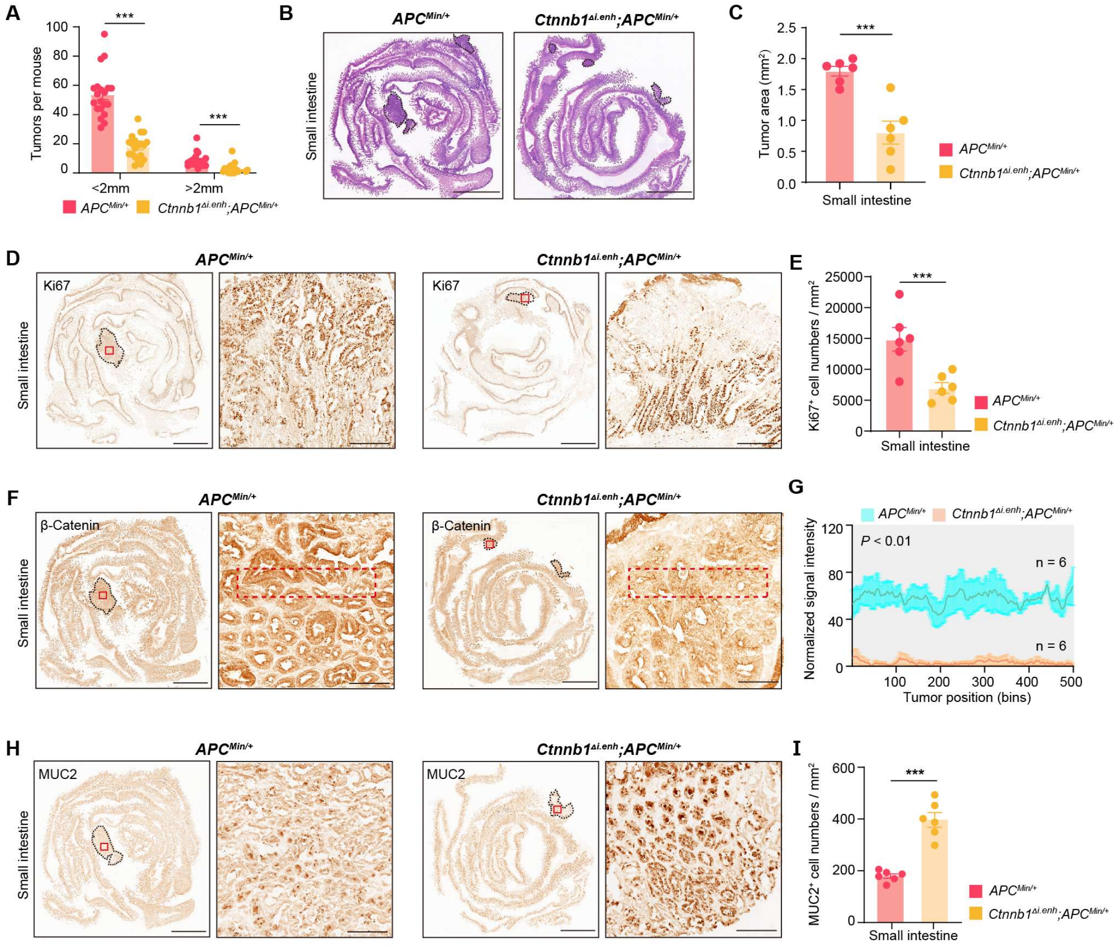


**Figure 3 - figure supplement 1. Knocking out ieCtnnb1 inhibits tumorigenesis of small intestine.** (A) Statistical analyses of tumor counts in small intestines of 5-month-old *Apc*$^{\text{Min/+}}$ (n = 20) and *Ctnnb1*$^{\Delta i.enh}$;*Apc*$^{\text{Min/+}}$ (n = 20) mice. (B) Representative H&E staining images of small intestine sections of 5-month-old *Apc*$^{\text{Min/+}}$ and *Ctnnb1*$^{\Delta i.enh}$;*Apc*$^{\text{Min/+}}$ mice. (C) The statistical analysis of small intestinal tumor area in 5-month-old *Apc*$^{\text{Min/+}}$ (n = 6) and *Ctnnb1*$^{\Delta i.enh}$;*Apc*$^{\text{Min/+}}$ (n = 6) mice. (D-E) Immunohistochemistry (D) and quantification (E) of Ki67+ cells in small intestinal tumors of 5-month-old *Apc*$^{\text{Min/+}}$ (n = 6) and *Ctnnb1*$^{\Delta i.enh}$;*Apc*$^{\text{Min/+}}$ (n = 6) mice. (F-G) Immunohistochemistry (F) and signal intensity statistics (G, red dashed boxes of F) of β-Catenin in small intestinal tumors of 5-month-old *Apc*$^{\text{Min/+}}$ (n = 6) and *Ctnnb1*$^{\Delta i.enh}$;*Apc*$^{\text{Min/+}}$ (n = 6) mice. (H-I) Immunohistochemistry (H) and quantification (I) of MUC2+ cells in small intestinal tumors of 5-month-old *Apc*$^{\text{Min/+}}$ (n = 6) and *Ctnnb1*$^{\Delta i.enh}$;*Apc*$^{\text{Min/+}}$ (n = 6) mice. Scale bars, 4 mm (B, D, F and H), 200 μm (magnified views in D and H), 100 μm (magnified views in F). Quantification data are shown as means

± SEM, statistical significance was determined using an unpaired two-tailed Student's *t*-test (C, E, G and I) or Multiple *t*-tests - one per row (A). *$P < 0.05$, **$P < 0.01$, ***$P < 0.001$, and ****$P < 0.0001$. ns, not significant.

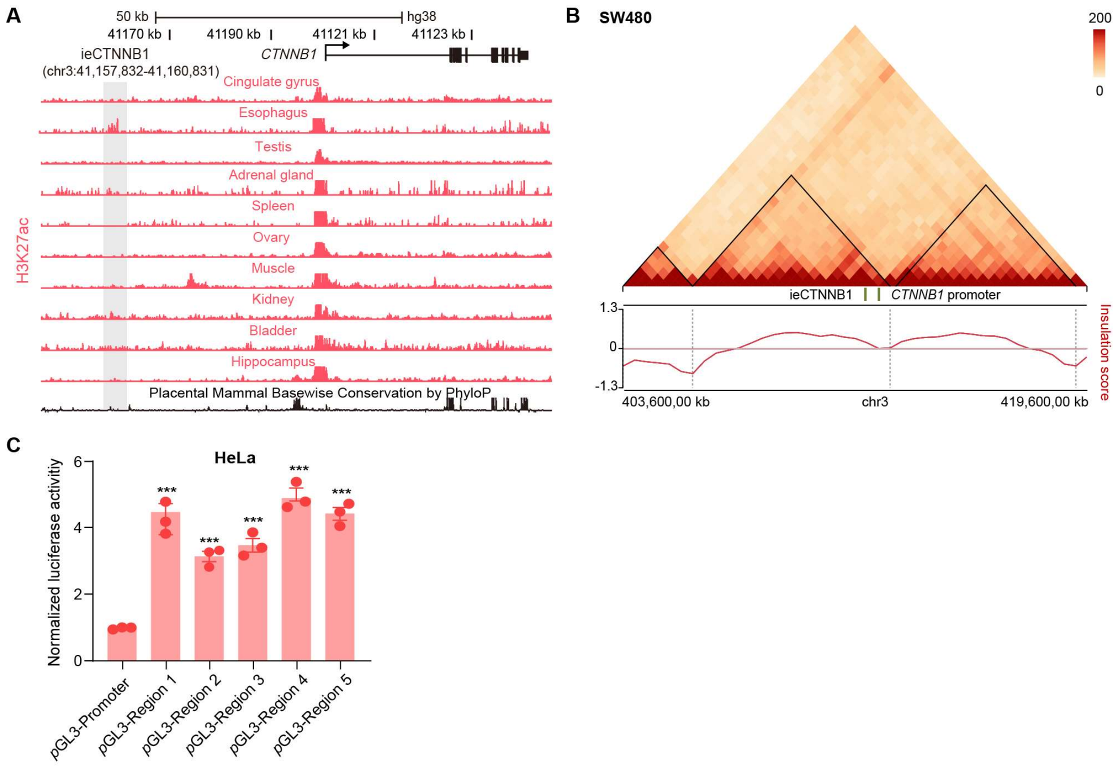


**Figure 4 - figure supplement 1. ieCTNNB1 is the intestinal enhancer of human *CTNNB1*.** (A) Schematic representation of the upstream region of human *CTNNB1* gene and the location of ieCTNNB1 (gray shading). Enrichment of H3K27ac ChIP-seq signals in indicated human tissues was shown. Data were obtained from ENCODE. (B) Hi-C data of SW480 cells. Boundaries of the TADs and locations of *CTNNB1* promoter and ieCTNNB1 (green bar) were marked. (C) Luciferase reporter assays of HeLa cells transfected with indicated plasmids for 48 hours. Quantification data are shown as means ± SEM, statistical significance was determined using one-way ANOVA analysis (C). $^*P < 0.05$, $^{**}P < 0.01$, $^{***}P < 0.001$, and $^{****}P < 0.0001$. ns, not significant.

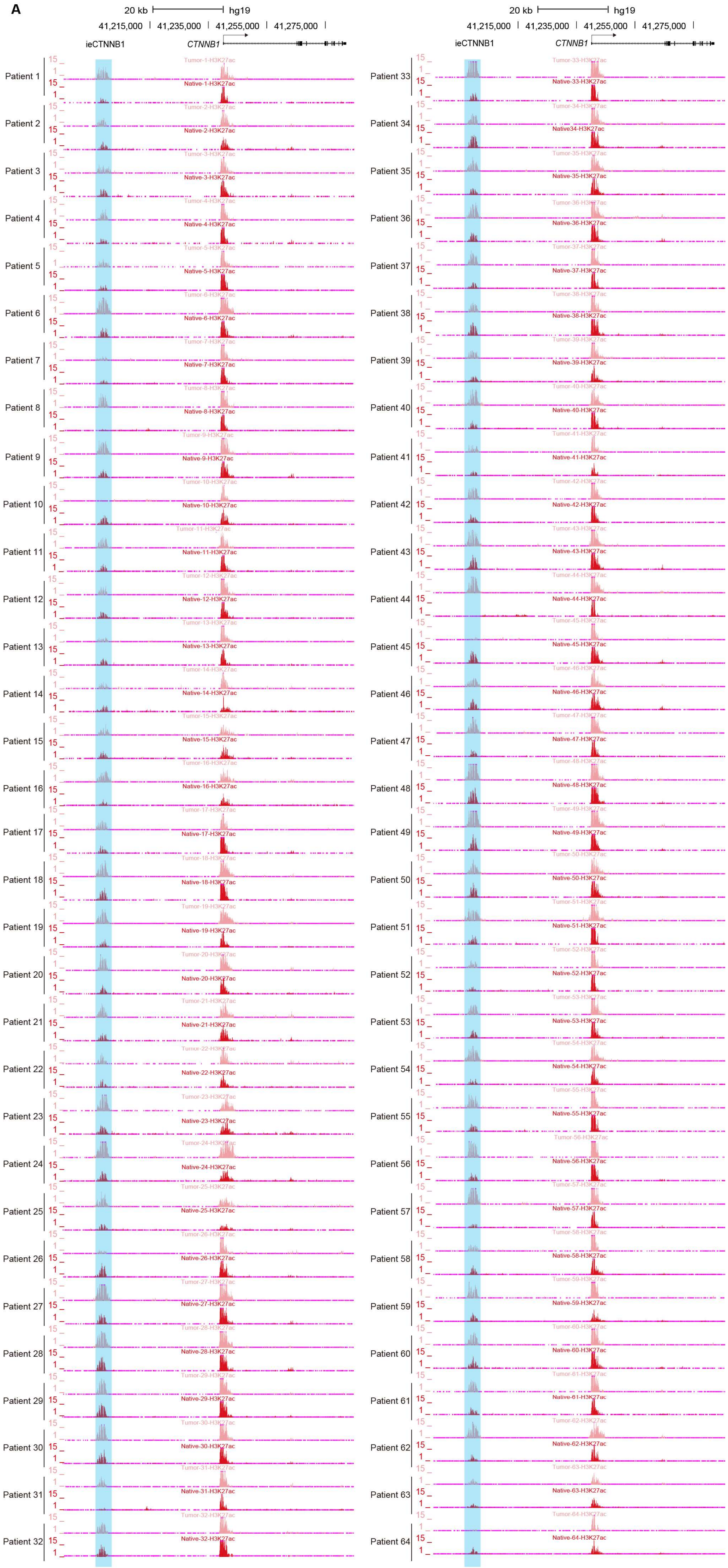

A
20 kb
hg19
41,215,000
41,235,000
41,255,000
41,275,000
ieCTNNB1
CTNNB1
Patient 1
Patient 2
Patient 3
Patient 4
Patient 5
Patient 6
Patient 7
Patient 8
Patient 9
Patient 10
Patient 11
Patient 12
Patient 13
Patient 14
Patient 15
Patient 16
Patient 17
Patient 18
Patient 19
Patient 20
Patient 21
Patient 22
Patient 23
Patient 24
Patient 25
Patient 26
Patient 27
Patient 28
Patient 29
Patient 30
Patient 31
Patient 32
20 kb
hg19
41,215,000
41,235,000
41,255,000
41,275,000
ieCTNNB1
CTNNB1
Patient 33
Patient 34
Patient 35
Patient 36
Patient 37
Patient 38
Patient 39
Patient 40
Patient 41
Patient 42
Patient 43
Patient 44
Patient 45
Patient 46
Patient 47
Patient 48
Patient 49
Patient 50
Patient 51
Patient 52
Patient 53
Patient 54
Patient 55
Patient 56
Patient 57
Patient 58
Patient 59
Patient 60
Patient 61
Patient 62
Patient 63
Patient 64

**Figure 5 - figure supplement 1. ieCTNNB1 is activated in colorectal cancer.**

(A) H3K27ac ChIP-seq signals at ieCTNNB1 (blue shading) in paired tumor (top) and native (bottom) tissues of patients with colorectal cancer (n = 64).

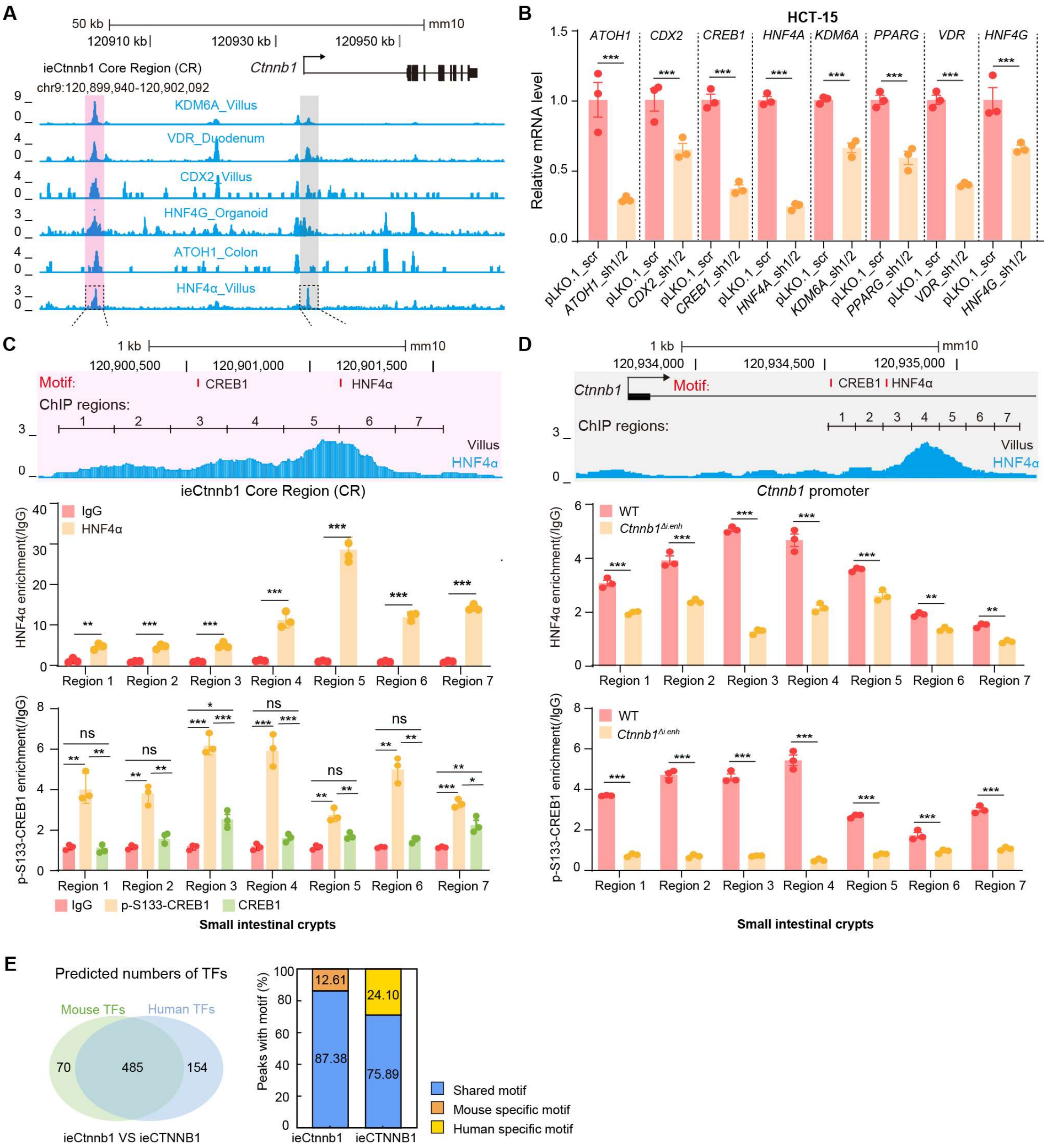


**Figure 6 - figure supplement 1. HNF4$\alpha$ and p-S133-CREB1 associate with ieCtnnb1 to regulate *Ctnnb1*'s transcription.** (A) ChIP-seq tracks of indicated trans-acting factors enriched at ieCtnnb1 (pink shading) and *Ctnnb1* promoter (grey shading) in mouse intestinal tissues and organoids. Shaded regions were enlarged in C (pink) and D (grey) respectively. (B) Relative mRNA levels of indicated genes in HCT-15 cells transfected with indicated shRNA-expressing plasmids for 48 hours. (C-D) Top: schematic diagram showing the enrichment of HNF4$\alpha$ at ieCtnnb1 (C) and *Ctnnb1*'s promoter (D). Locations of the HNF4$\alpha$ and CREB1 binding motif sites were indicated. Middle and bottom: ChIP-qPCR showing enrichment of HNF4$\alpha$ (middle), CREB1 and p-S133-

CREB1 (bottom) at ieCtnnb1 (C) and *Ctnnb1*'s promoter (D) in small intestinal crypts. ChIP regions were indicated. (E) The comparison of numbers (left) and proportions (right) of species-specific and shared transcription factor binding sites within ieCtnnb1 and ieCTNNB1. Quantification data are shown as means ± SEM, statistical significance was determined using an unpaired two-tailed Student's *t*-test (B) and Multiple *t*-tests - one per row (C and D). *$P < 0.05$, **$P < 0.01$, ***$P < 0.001$, and ****$P < 0.0001$. ns, not significant.

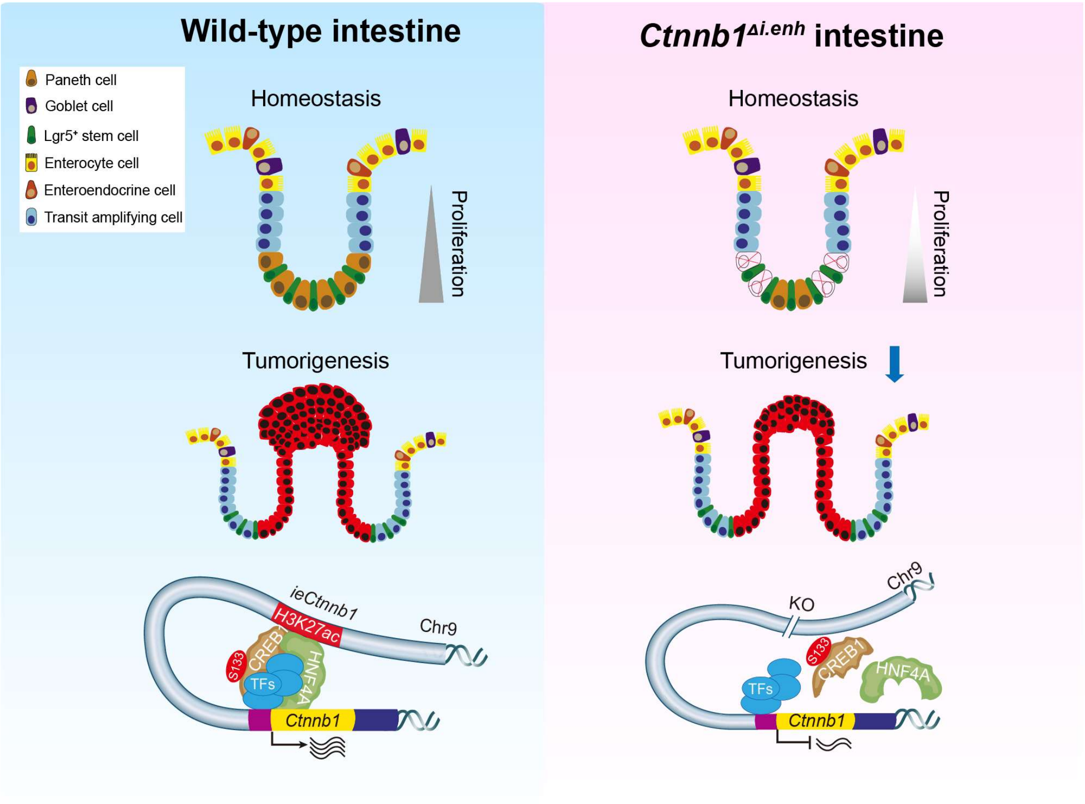


**Figure supplement 8. The working model.** ieCtnnb1, the intestinal enhancer of *Ctnnb1*, balances epithelial homeostasis and tumorigenesis by transcriptionally controlling Wnt signaling dosage.

**Table S1. Summary of high throughput data used in this study.**

| Tissue | Species | Age | Strategy | Antibody | Serial number |
|---|---|---|---|---|---|
| Intestine | Mus musculus | E14.5 | ChIP-seq | H3K27ac | ENCSR424END |
| Intestine | Mus musculus | E15.5 | ChIP-seq | H3K27ac | ENCSR599GVS |
| Intestine | Mus musculus | E16.5 | ChIP-seq | H3K27ac | ENCSR639DND |
| Intestine | Mus musculus | P0 | ChIP-seq | H3K27ac | ENCSR642VYW |
| Small Intestine | Mus musculus | 2 months | ChIP-seq | H3K27ac | ENCSR000CCQ |
| Intestine | Mus musculus | E14.5 | ChIP-seq | H3K4me3 | ENCSR464MQU |
| Intestine | Mus musculus | E15.5 | ChIP-seq | H3K4me3 | ENCSR410YIY |
| Intestine | Mus musculus | E16.5 | ChIP-seq | H3K4me3 | ENCSR572KYR |
| Intestine | Mus musculus | P0 | ChIP-seq | H3K4me3 | ENCSR198ACZ |
| Small Intestine | Mus musculus | 2 months | ChIP-seq | H3K4me3 | ENCSR000CCS |
| Intestine | Mus musculus | E14.5 | ChIP-seq | H3K4me1 | ENCSR157LYR |
| Intestine | Mus musculus | E15.5 | ChIP-seq | H3K4me1 | ENCSR051CUH |
| Intestine | Mus musculus | E16.5 | ChIP-seq | H3K4me1 | ENCSR829YGD |
| Intestine | Mus musculus | P0 | ChIP-seq | H3K4me1 | ENCSR159RVN |
| Small Intestine | Mus musculus | 2 months | ChIP-seq | H3K4me1 | ENCSR000CCR |
| Intestine | Mus musculus | E14.5 | DNase-seq | / | ENCSR655WKX |
| Large Intestine | Mus musculus | 2 months | DNase-seq | / | ENCSR000CNH |
| Intestine | Mus musculus | E14.5 | ChIP-seq | H3k36me3 | ENCSR953KTY |
| Intestine | Mus musculus | E15.5 | ChIP-seq | H3k36me3 | ENCSR919DDC |
| Intestine | Mus musculus | E16.5 | ChIP-seq | H3k36me3 | ENCSR272XPJ |
| Intestine | Mus musculus | P0 | ChIP-seq | H3k36me3 | ENCSR483KOD |
| Small Intestine | Mus musculus | 2 months | ChIP-seq | H3k36me3 | ENCSR000CFS |
| Stomach | Mus musculus | E14.5 | ChIP-seq | H3K27ac | ENCSR316CNR |
| Stomach | Mus musculus | E15.5 | ChIP-seq | H3K27ac | ENCSR929SEW |
| Stomach | Mus musculus | E16.5 | ChIP-seq | H3K27ac | ENCSR546ANT |
| Stomach | Mus musculus | P0 | ChIP-seq | H3K27ac | ENCSR346FJG |
| Stomach | Mus musculus | E15.5 | ChIP-seq | H3K4me1 | ENCSR548BKP |
| Stomach | Mus musculus | E14.5 | ChIP-seq | H3K4me1 | ENCSR335WME |
| Stomach | Mus musculus | E16.5 | ChIP-seq | H3K4me1 | ENCSR907CPZ |
| Stomach | Mus musculus | P0 | ChIP-seq | H3K4me1 | ENCSR940CMI |
| Stomach | Mus musculus | E15.5 | ChIP-seq | H3K4me3 | ENCSR522LXN |
| Stomach | Mus musculus | E14.5 | ChIP-seq | H3K4me3 | ENCSR023VJO |
| Stomach | Mus musculus | E16.5 | ChIP-seq | H3K4me3 | ENCSR684UWM |
| Stomach | Mus musculus | P0 | ChIP-seq | H3K4me3 | ENCSR916CBN |
| Stomach | Mus musculus | P0 | DNase-seq | / | ENCSR969OPE |
| Stomach | Mus musculus | E14.5 | ChIP-seq | H3k36me3 | ENCSR581FAT |
| Stomach | Mus musculus | E15.5 | ChIP-seq | H3k36me3 | ENCSR599PKR |
| Stomach | Mus musculus | E16.5 | ChIP-seq | H3k36me3 | ENCSR872WGX |
| Stomach | Mus musculus | P0 | ChIP-seq | H3k36me3 | ENCSR516KLO |
| Liver | Mus musculus | E14.5 | ChIP-seq | H3K27ac | ENCSR075SNV |
| Liver | Mus musculus | E15.5 | ChIP-seq | H3K27ac | ENCSR479LFP |

| | | | | | |
|---|---|---|---|---|---|
| Liver | Mus musculus | E16.5 | ChIP-seq | H3K27ac | ENCSR802RET |
| Liver | Mus musculus | P0 | ChIP-seq | H3K27ac | ENCSR616TJM |
| Liver | Mus musculus | 2 months | ChIP-seq | H3K27ac | ENCSR000CDH |
| Liver | Mus musculus | E14.5 | ChIP-seq | H3K4me3 | ENCSR433ESG |
| Liver | Mus musculus | E15.5 | ChIP-seq | H3K4me3 | ENCSR577SDJ |
| Liver | Mus musculus | E16.5 | ChIP-seq | H3K4me3 | ENCSR252GKD |
| Liver | Mus musculus | P0 | ChIP-seq | H3K4me3 | ENCSR653AVN |
| Liver | Mus musculus | 2 months | ChIP-seq | H3K4me3 | ENCSR000CAP |
| Liver | Mus musculus | E14.5 | ChIP-seq | H3K4me1 | ENCSR000CDW |
| Liver | Mus musculus | E15.5 | ChIP-seq | H3K4me1 | ENCSR133EGP |
| Liver | Mus musculus | E16.5 | ChIP-seq | H3K4me1 | ENCSR487OLC |
| Liver | Mus musculus | P0 | ChIP-seq | H3K4me1 | ENCSR308GFM |
| Liver | Mus musculus | 2 months | ChIP-seq | H3K4me1 | ENCSR000CAO |
| Liver | Mus musculus | E14.5 | DNase-seq | / | ENCSR000CNJ |
| Liver | Mus musculus | E14.5 | ChIP-seq | H3k36me3 | ENCSR670YXP |
| Liver | Mus musculus | E15.5 | ChIP-seq | H3k36me4 | ENCSR510CGB |
| Liver | Mus musculus | E16.5 | ChIP-seq | H3k36me5 | ENCSR569DBO |
| Liver | Mus musculus | P0 | ChIP-seq | H3k36me6 | ENCSR656AMS |
| Liver | Mus musculus | 2 months | ChIP-seq | H3k36me7 | ENCSR000CEO |
| Forebrain | Mus musculus | P0 | ChIP-seq | H3K27ac | ENCSR094TTT |
| Heart | Mus musculus | 2 months | ChIP-seq | H3K27ac | ENCSR000CDF |
| Lung | Mus musculus | P0 | ChIP-seq | H3K27ac | ENCSR884MYD |
| Limb | Mus musculus | E15.5 | ChIP-seq | H3K27ac | ENCSR988BRP |
| Craniofacial | Mus musculus | E14.5 | ChIP-seq | H3K27ac | ENCSR481SGM |
| Cortical plate | Mus musculus | 2 months | ChIP-seq | H3K27ac | ENCSR000CDD |
| Midbrain | Mus musculus | P0 | ChIP-seq | H3K27ac | ENCSR672ZXY |
| Hindbrain | Mus musculus | P0 | ChIP-seq | H3K27ac | ENCSR332JYZ |
| Testis | Mus musculus | 2 months | ChIP-seq | H3K27ac | ENCSR000CCU |
| Cerebellum | Mus musculus | 2 months | ChIP-seq | H3K27ac | ENCSR000CDC |
| Gastrocnemius | Mus musculus | 2 months | ChIP-seq | H3K27ac | ENCSR714ZJT |
| Kidney | Mus musculus | 2 months | ChIP-seq | H3K27ac | ENCSR000CDG |
| Small Intestine | Homo sapiens | 30 years | ChIP-seq | H3K27ac | ENCSR655XLM |
| Sigmoid colon | Homo sapiens | 34 years | ChIP-seq | H3K27ac | ENCSR561YSH |
| Small Intestine | Homo sapiens | 30 years | ChIP-seq | H3K4me3 | ENCSR944QSH |
| Sigmoid colon | Homo sapiens | 34 years | ChIP-seq | H3K4me3 | ENCSR792IJA |
| Small Intestine | Homo sapiens | 30 years | ChIP-seq | H3K4me1 | ENCSR538JMW |
| Sigmoid colon | Homo sapiens | 34 years | ChIP-seq | H3K4me1 | ENCSR782OZZ |
| Small Intestine | Homo sapiens | 30 years | ChIP-seq | H3K36me3 | ENCSR073YZL |
| Sigmoid colon | Homo sapiens | 34 years | ChIP-seq | H3K36me3 | ENCSR445RFF |
| Small Intestine | Homo sapiens | 34 years | DNase-seq | / | ENCSR931UQB |
| Sigmoid colon | Homo sapiens | 37 years | DNase-seq | / | ENCSR923JYH |

| | | | | | |
|---|---|---|---|---|---|
| Large intestine | Mus musculus | 2 months | DNase-seq | / | ENCSR000CNH |
| Hippocampus | Homo sapiens | 73 years | ChIP-seq | H3K27ac | ENCSR321LKT |
| Kidney | Homo sapiens | 50 years | ChIP-seq | H3K27ac | ENCSR438SPO |
| Urinary bladder | Homo sapiens | 34 years | ChIP-seq | H3K27ac | ENCSR054BKO |
| Muscle of leg | Homo sapiens | 110 days | ChIP-seq | H3K27ac | ENCSR687ZCM |
| Cingulate gyrus | Homo sapiens | 75 years | ChIP-seq | H3K27ac | ENCSR604JDV |
| Spleen | Homo sapiens | 30 years | ChIP-seq | H3K27ac | ENCSR086XCT |
| Ovary | Homo sapiens | 30 years | ChIP-seq | H3K27ac | ENCSR268JQE |
| Testis | Homo sapiens | 37 years | ChIP-seq | H3K27ac | ENCSR136ZQZ |
| Adrenal gland | Homo sapiens | 30 years | ChIP-seq | H3K27ac | ENCSR642HHF |
| Esophagus | Homo sapiens | 30 years | ChIP-seq | H3K27ac | ENCSR645SYH |
| Small Intestine | Mus musculus | Adult | ChIP-seq | HNF4$\alpha$ | GSM851120 |
| Organoids | Mus musculus | / | ChIP-seq | HNF4G | GSM3132969 |
| Colon | Mus musculus | Adult | ChIP-seq | ATOH1 | GSM2185705 |
| Intestinal villus | Mus musculus | Adult | ChIP-seq | KDM6A | GSM2610642 |
| Intestinal villus | Mus musculus | Adult | ChIP-seq | CDX2 | GSM2610627 |
| Duodenum | Mus musculus | Adult | ChIP-seq | VDR | GSM1694861 |
| Caco-2 | Homo sapiens | / | ChIP-seq | HNF4$\alpha$ | GSM575229 |
| LS180 | Homo sapiens | / | ChIP-seq | CDX2 | GSM791413 |
| HT29 | Homo sapiens | / | ChIP-seq | PPARG | GSM2042856 |
| LoVo | Homo sapiens | / | ChIP-seq | CREB1 | GSM1239450 |

**Table S2. Primers used in this study.**

| Primers | Sequence (5'-3') | Description |
|---|---|---|
| shRNA-*ATOH1*-1 | cccttccagcaaacaggtgaa | used for RT-qPCR |
| shRNA-*ATOH1*-2 | agttgccggactcgcttctca | used for RT-qPCR |
| shRNA-*CDX2*-1 | caaatatcgagtggtgtacac | used for RT-qPCR |
| shRNA-*CDX2*-2 | acgtgagcatgtaccctagct | used for RT-qPCR |
| shRNA-*CREB*-1 | acggtgccaactccaatttac | used for RT-qPCR |
| shRNA-*CREB*-2 | acagcacccactagcactatt | used for RT-qPCR |
| shRNA-*PPARG*-1 | ctggcctccttgatgaataaa | used for RT-qPCR |
| shRNA-*PPARG*-2 | gacaacagacaaatcaccatt | used for RT-qPCR |
| shRNA-*VDR*-1 | ttggctttgctaagatgatac | used for RT-qPCR |
| shRNA-*VDR*-2 | cctccagttcgtgtgaatgat | used for RT-qPCR |
| shRNA-*HNF4A*-1 | tcacctgatgcaggaacatat | used for RT-qPCR |
| shRNA-*HNF4A*-2 | tcttcggcatggccaagattg | used for RT-qPCR |
| shRNA-*HNF4G*-1 | gtgacagaataagcaccagaa | used for RT-qPCR |
| shRNA-*HNF4G*-2 | cgtggcaaatgattgagcaaa | used for RT-qPCR |
| shRNA-*KDM6A*-1 | ccgcgcaaatagaaataattt | used for RT-qPCR |
| shRNA-*KDM6A*-2 | gcagcacgaattaagtattta | used for RT-qPCR |
| *ATOH1*-F | ccttccagcaaacaggtgaat | used for RT-qPCR |
| *ATOH1*-R | ttgttgaacgacgggataaca | used for RT-qPCR |
| *CREB*-F | attcacaggagtcagtggatagt | used for RT-qPCR |
| *CREB*-R | caccgttacagtggtgatgg | used for RT-qPCR |
| *PPARG*-F | gatgccagcgactttgactc | used for RT-qPCR |
| *PPARG*-R | acccacgtcatcttcaggga | used for RT-qPCR |
| *CDX2*-F | gacgtgagcatgtaccctagc | used for RT-qPCR |
| *CDX2*-R | gcgtagccattccagtcct | used for RT-qPCR |
| *VDR*-F | gtggacatcggcatgatgaag | used for RT-qPCR |
| *VDR*-R | ggtcgtaggtcttatggtggg | used for RT-qPCR |
| *HNF4A*-F | cacgggcaaacactacggt | used for RT-qPCR |
| *HNF4A*-R | ttgaccttcgagtgctgatcc | used for RT-qPCR |
| *HNF4G*-F | ttgcaggttcagtcggcaat | used for RT-qPCR |
| *HNF4G*-R | tttcattcccgctctaaaacact | used for RT-qPCR |
| *KDM6A*-F | ttcctcggaaggtgctattca | used for RT-qPCR |
| *KDM6A*-R | gaggctggttgcaggattca | used for RT-qPCR |
| *Ctnnb1*-F | atggagccggacagaaaagc | used for RT-qPCR |
| *Ctnnb1*-R | cttgccactcagggaagga | used for RT-qPCR |
| *CTNNB1*-F | aaagcggctgttagtcactgg | used for RT-qPCR |
| *CTNNB1*-R | cgagtcattgcatactgtccat | used for RT-qPCR |
| *Gapdh*-F | aggtcggtgtgaacggatttg | used for RT-qPCR |
| *Gapdh*-R | tgtagaccatgtagttgaggtca | used for RT-qPCR |
| *ACTB*-F | agtgcgacgtggacatccg | used for RT-qPCR |
| *ACTB*-R | tggctctaacagtccgcctag | used for RT-qPCR |

| | | |
|---|---|---|
| ieCtnnb1-ChIP-1F | tgactaaggacaggcctttcccc | used for ChIP-qPCR |
| ieCtnnb1-ChIP-1R | ggcagtcacaggtcaggtctcat | used for ChIP-qPCR |
| ieCtnnb1-ChIP-2F | gcagaggtaccagagcaaaggtg | used for ChIP-qPCR |
| ieCtnnb1-ChIP-2R | gctggaatccttgaagacctgct | used for ChIP-qPCR |
| ieCtnnb1-ChIP-3F | ctgacgtcttgacattcctgttga | used for ChIP-qPCR |
| ieCtnnb1-ChIP-3R | gcctgaacgatggagattacatcc | used for ChIP-qPCR |
| ieCtnnb1-ChIP-4F | cctcctgagagttctgccatcct | used for ChIP-qPCR |
| ieCtnnb1-ChIP-4R | gcttgaacaggcagactttgtaacc | used for ChIP-qPCR |
| ieCtnnb1-ChIP-5F | tttgtggcaagtcttggacagct | used for ChIP-qPCR |
| ieCtnnb1-ChIP-5R | tggttctgccttgtgatcagcaa | used for ChIP-qPCR |
| ieCtnnb1-ChIP-6F | ctgtcagagggcaaagccgttta | used for ChIP-qPCR |
| ieCtnnb1-ChIP-6R | tgtgggaatcgggaagctctg | used for ChIP-qPCR |
| ieCtnnb1-ChIP-7F | gcttctttccgggggcaaagt | used for ChIP-qPCR |
| ieCtnnb1-ChIP-7R | gggaatgtcaggcacagtgca | used for ChIP-qPCR |
| *pCtnnb1*-ChIP-1F | gcagatgtctcagtgcagtctct | used for ChIP-qPCR |
| *pCtnnb1*-ChIP-1R | ccgaaatacaaaggccacaagct | used for ChIP-qPCR |
| *pCtnnb1*-ChIP-2F | gcttgtggcctttgtatttcggt | used for ChIP-qPCR |
| *pCtnnb1*-ChIP-2R | cggggatttctgaatgtttaaggtga | used for ChIP-qPCR |
| *pCtnnb1*-ChIP-3F | ccccgaaattaaaatgaagtgcctc | used for ChIP-qPCR |
| *pCtnnb1*-ChIP-3R | ggacaaaggttaggtaggtcacaga | used for ChIP-qPCR |
| *pCtnnb1*-ChIP-4F | acctttgtcctcaaggccgag | used for ChIP-qPCR |
| *pCtnnb1*-ChIP-4R | gcctgggagggtttgttgtagag | used for ChIP-qPCR |
| *pCtnnb1*-ChIP-5F | caaaccctcccaggctgaagtc | used for ChIP-qPCR |
| *pCtnnb1*-ChIP-5R | ttacagactgtgaggcccagga | used for ChIP-qPCR |
| *pCtnnb1*-ChIP-6F | gggaaacatttaatcgaatgcaggcg | used for ChIP-qPCR |
| *pCtnnb1*-ChIP-6R | tcaggaagactgactgagaagcac | used for ChIP-qPCR |
| *pCtnnb1*-ChIP-7F | caactgttatggtgaccccaacc | used for ChIP-qPCR |
| *pCtnnb1*-ChIP-7R | ggtcacctaagaccgtgggaaaa | used for ChIP-qPCR |
| ieCTNNB1-ChIP-1F | tttctagggcatctgctgcaaag | used for ChIP-qPCR |
| ieCTNNB1-ChIP-1R | gacaaactataagtggccattgacca | used for ChIP-qPCR |
| ieCTNNB1-ChIP-2F | cacagacagttctattaaacaacact | used for ChIP-qPCR |
| ieCTNNB1-ChIP-2R | gtgagagtggaaggtggattaaaaa | used for ChIP-qPCR |
| ieCTNNB1-ChIP-3F | ggggatgttgtaggtgctcaata | used for ChIP-qPCR |
| ieCTNNB1-ChIP-3R | gaataaatcaggcccacttccaa | used for ChIP-qPCR |
| ieCTNNB1-ChIP-4F | tttggtagctgattgagagcttg | used for ChIP-qPCR |
| ieCTNNB1-ChIP-4R | gctcccataaattatgtcaccattc | used for ChIP-qPCR |
| ieCTNNB1-ChIP-5F | gtcatacctacccattacctcttc | used for ChIP-qPCR |
| ieCTNNB1-ChIP-5R | taaagagacacttgtcctcatgg | used for ChIP-qPCR |
| ieCTNNB1-ChIP-6F | tcctggagttcctgtaacccagtt | used for ChIP-qPCR |
| ieCTNNB1-ChIP-6R | tactgcttccaactggctgtgtg | used for ChIP-qPCR |
| *pCTNNB1*-ChIP-1F | gctgcttaatcgatagctttctc | used for ChIP-qPCR |
| *pCTNNB1*-ChIP-1R | ccacttgtcactaggtatcaatag | used for ChIP-qPCR |

| | | |
|---|---|---|
| *pCTNNB1*-ChIP-2F | gcgctctggagctaatccatttc | used for ChIP-qPCR |
| *pCTNNB1*-ChIP-2R | aaggctgtgaactctccgtagaa | used for ChIP-qPCR |
| *pCTNNB1*-ChIP-3F | gctgaacagcctgctgagaggt | used for ChIP-qPCR |
| *pCTNNB1*-ChIP-3R | ttgtggtctgtgccgcacactc | used for ChIP-qPCR |
| *pCTNNB1*-ChIP-4F | gagatgccaccttccgcagg | used for ChIP-qPCR |
| *pCTNNB1*-ChIP-4R | aaggtggccctggtatcctc | used for ChIP-qPCR |
| *pCTNNB1*-ChIP-5F | atgcagaccacagcgccctca | used for ChIP-qPCR |
| *pCTNNB1*-ChIP-5R | agcagtctgctgccgtctgag | used for ChIP-qPCR |
| *pCTNNB1*-ChIP-6F | ggtctgaggagcagcttcagtc | used for ChIP-qPCR |
| *pCTNNB1*-ChIP-6R | ataaggaaaggagcgcccaagc | used for ChIP-qPCR |
| sg1 | tgactccaactacaagcgag | used for CRISPRa andCRISPRi |
| sg2 | ttacgagctgccaactctca | used for CRISPRa andCRISPRi |
| sg3 | ctgtctaggtaggcggtagt | used for CRISPRa andCRISPRi |
| *Ctnnb1*$^{\Delta i.enh}$ -WT-F | gtcctgtccgtcactattatcctggc | used for validation of *Ctnnb1*$^{\Delta i.enh}$ mice |
| *Ctnnb1*$^{\Delta i.enh}$ -WT-R | ccactgccctgctaaagcattggt | used for validation of *Ctnnb1*$^{\Delta i.enh}$ mice |
| *Ctnnb1*$^{\Delta i.enh}$ -Mut-R | tgattagttctccgggaagcccagt | used for validation of *Ctnnb1*$^{\Delta i.enh}$ mice |
| *APC*$^{Min/+}$-F | atctcatggcaaacagacct | used for validation of *APC*$^{Min/+}$mice |
| *APC*$^{Min/+}$ -R | tcacaaatcatctcgcaga | used for validation of *APC*$^{Min/+}$mice |
| *Lgr5-EGFP*-WT-F | ctgctctctgctcccagtct | used for validation of Lgr5-EGFP mice |
| *Lgr5-EGFP*-WT-R | ataccccatccctttgagc | used for validation of Lgr5-EGFP mice |
| *Lgr5-EGFP*-Mut-R | gaacttcagggtcagcttgc | used for validation of Lgr5-EGFP mice |
| *LacZ* F | atcctctgcatggtcaggtc | used for validation of *H11*$^{i.enh}$/ *H11*$^{hi.enh}$ and BAT-Gal mice |
| *LacZ* R | cgtggcctgattcattcc | used for validation of *H11*$^{i.enh}$/ *H11*$^{hi.enh}$ and BAT-Gal mice |
| sgRNA-1 | tgttggctcagcagacacccagg | used for construction of *H11*$^{i.enh}$ mice |
| sgRNA-2 | actgcctcctcagcttcaagagg | used for construction of *H11*$^{i.enh}$ mice |
| EGE-WL-020-T7-sgRNA3 | ctatttctagctctaaaactttgggg atcaagtaaggggcctatagtgagt | used for construction of *Ctnnb1*$^{\Delta i.enh}$ mice |
| EGE-WL-020-T7-sgRNA13 | ctatttctagctctaaaaccctcatca taggaaggcttcctatagtgagtcgt | used for construction of *Ctnnb1*$^{\Delta i.enh}$ mice |

**Table S3. HNF4$\alpha$ and CREB1 motifs analyses at ieCtnnb1/ieCTNNB1 and *Ctnnb1/CTNNB1* promoter.**

| | Name | Matrix ID | Sequence logo | Score | Relative score | Predicted sequence |
|---|---|---|---|---|---|---|
| ieCtnnb1 (*Mus musculus*) | CREB1 | MA0018.2 | 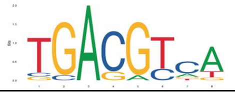 | 9.92 | 0.93 | TGACGTCT |
| | HNF4$\alpha$ | MA0114.3 | 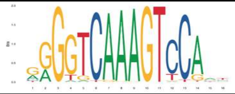 | 16.39 | 0.90 | GGGGGCAAA GTCTATC |
| *Ctnnb1* Promoter (*Mus musculus*) | CREB1 | MA0018.2 | 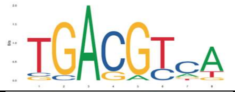 | 8.00 | 0.86 | TGAAGTCA |
| | HNF4$\alpha$ | MA0114.3 | 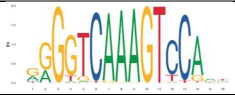 | 5.18 | 0.79 | CGGTTCCAA GTCTGGT |
| ieCTNNB1 *(Homo sapiens)* | CREB1 | MA0018.5 | 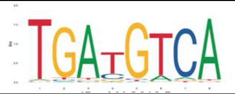 | 14.63 | 0.99 | TGATGTCA |
| | | MA0018.4 | 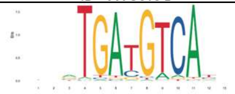 | 14.29 | 0.95 | CTCTGATGT CACC |
| | HNF4$\alpha$ | MA0114.2 | 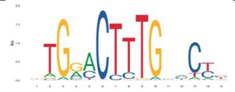 | 10.15 | 0.87 | CTAAACTGT GAACTC |
| | | MA0114.5 | 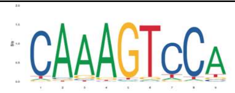 | 9.78 | 0.86 | CAGAGTCCT |
| ieCTNNB1 Promoter *(Homo sapiens)* | CREB1 | MA0018.1 | 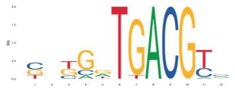 | 10.00 | 0.89 | CGGGGTGAC GGC |
| | HNF4$\alpha$ | MA0114.2 | 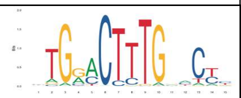 | 14.99 | 0.93 | GAGGACTTT GAACCG |
